# Sound-evoked responses of distinct neuron classes from the tail of the striatum

**DOI:** 10.1101/2022.02.26.482132

**Authors:** Matthew B. Nardoci, Anna A. Lakunina, Devin C. Henderling, Jewlyssa C. Pedregon, Santiago Jaramillo

## Abstract

Given its inputs from auditory structures and neuromodulatory systems, the posterior tail of the striatum is ideally positioned to influence behavioral responses to acoustic stimuli according to context and previous rewards. Results from previous studies indicate that neurons in this striatal region display selective responses to sounds. However, it is not clear whether different striatal cell classes code for distinct features of sounds, nor how different striatal output pathways may use acoustic information to guide behavior. Here we compared the sound-evoked responses of posterior striatal neurons that form the striatal direct pathway (and express the dopamine receptor D1) to the responses of neighboring neurons in naive mice. We achieved this via optogenetic photo-identification of D1-expressing neurons during extracellular electrophysiological recordings in awake head-fixed mice of both sexes. We found that the frequency tuning of sound-responsive direct-pathway striatal neurons is comparable to that of their sound-responsive neighbors. Moreover, we found that both populations encode amplitude modulated sounds in a similar fashion. These results suggest that different classes of neurons in the posterior striatum of naive animals have similar access to acoustic features conveyed by the auditory system even outside the context of an auditory task.

## Introduction

The striatum, as the primary input structure of the basal ganglia and a target of extensive dopaminergic inputs, is ideally positioned to influence behavioral responses to sensory stimuli according to context and previous rewards. Neurons in the posterior tail of the striatum receive numerous inputs from the auditory thalamus and the auditory cortex (Hintiryan et al., 2016; Ponvert and Jaramillo, 2019; Chen et al., 2019; Valjent and Gangarossa, 2021), and these striatal neurons display robust responses to sounds (Bordi and LeDoux, 1992; Znamenskiy and Zador, 2013; Zhong et al., 2014; Guo et al., 2018). It is not known, however, whether different striatal cell classes code for distinct features of sounds, nor how different striatal output pathways may use acoustic information to guide behavior. A key step toward understanding the processing of sounds by striatal circuits and the role of different striatal cells in auditory learning is the characterization of sound evoked responses by distinct striatal neuron classes in naive animals.

The striatum, including the posterior tail portion, is composed of a range of neuron classes with different gene expression and physiological profiles, including fast-spiking parvalbumin-expressing interneurons, spontaneously active cholinergic interneurons, and the abundant principal projection neurons (Kawaguchi, 1997; Valjent and Gangarossa, 2021). The large majority of these cells are medium spiny neurons that form the two main outputs of the striatum: the direct (striatonigral) pathway, composed of cells that express the dopamine receptor D1, and the indirect (striatopallidal) pathway, with cells that express the dopamine receptor D2 (Gerfen et al., 1990; Kreitzer and Malenka, 2008). Anatomical data suggest that for some striatal regions, cortical sensory neurons differentially innervate each of these striatal pathways (Lei et al., 2004; Wall et al., 2013). Moreover, excitatory synapses onto direct and indirect pathway neurons exhibit different synaptic transmission and plasticity properties (Kreitzer and Malenka, 2007). Together, these observations raise the possibility that neurons from different classes in the posterior striatum are differentially influenced by sensory signals.

Using optogenetic photo-identification of specific cell populations in the posterior tail of the striatum of naive mice, we characterized the sound-evoked responses of neurons that express the dopamine receptor D1 and compared these responses to those from neighboring neurons. We found that on average, sound-responsive D1-expressing posterior striatal neurons have similar frequency tuning to their non-D1 sound-responsive neighbors, and that both populations represent amplitude modulated noise in a similar fashion. These results suggest that different classes on neurons in the posterior striatum, likely including both major output pathways, have access to acoustic features conveyed from the auditory system in naive animals even outside the context of an auditory task.

## Materials and Methods

### Animals

A total of 15 transgenic adult DRD1::ChR2 mice of both sexes, were used in this study. This mouse line was generated by crossing animals that express Cre recombinase in neurons positive for the dopamine receptor D1 (RRID:MMRRC_036916-UCD) with mice that express the light-gated ion channel Channelrhodopsin-2 (ChR2) in a Cre-dependent manner (JAX 012569). All procedures were carried out in accordance with the National Institutes of Health standards and were approved by the University of Oregon Institutional Animal Care and Use Committee.

### Auditory stimuli

Experiments were performed inside a single-walled sound-isolation box (IAC Acoustics, North Aurora, IL). Auditory stimuli were presented in an open-field configuration from a speaker (MF1, TuckerDavis Technologies, Alachua, FL) contralateral to the side of electrophysiological recordings. Speakers were calibrated using an ultrasonic microphone (ANL-940-1, Med Associates, Inc., Fairfax, VT) to obtain the desired sound intensity level for frequencies between 1 kHz and 40 kHz. Stimuli were generated using Python software developed in-house (https://taskontrol.readthedocs.io/). The ensemble of auditory stimuli for evaluating frequency tuning consisted of pure-tone pips (100 ms duration) at 16 frequencies logarithmically-spaced between 2 kHz and 40 kHz and at 11 different intensities (15-70 dB SPL in 5 dB steps). We presented at least 9 repetitions per frequency-intensity combination with interstimulus intervals randomized in the range 0.7 − 0.9 seconds. Stimuli for evaluating responses to temporal sound features were sinusoidally amplitude modulated white noise at 11 modulation rates logarithmically spaced between 4 and 128 Hz (100% modulation depth, 500 ms duration, 60 dB SPL max). We presented at least 20 repetitions per condition using an interstimulus intervals randomized in the range 0.9 − 1.1 seconds. All stimuli had a 2 ms ramp up and ramp down. During sound presentation, mice were awake and head-fixed on top of a freely-moving wheel, leaving them free to move their limbs while their heads remained stationary.

### Surgical procedure

Mice were surgically implanted with a head-bar to allow for head-fixed recordings. Animals were anesthetized with isoflurane through a nose-cone on a stereotaxic apparatus. Bilateral craniotomies (AP: -1 mm to -2 mm from bregma, ML: ±2.9 mm to 4 mm from midline) and durotomies were performed to allow for acute recordings from the most posterior region of the dorsal striatum. Plastic wells were attached around each craniotomy and filled with a silicone elastomer (Sylgard 170, Dow-Corning) to protect the surface of the brain and retain moisture when not recording. All animals were monitored after surgery and recovered fully before electrophysiological experiments.

### Electrophysiological recordings and optogenetic stimulation

Electrical signals were collected with an RHD2000 acquisition system (Intan Technologies, Los Angeles, CA) and OpenEphys software (www.open-ephys.org), using 32-channel silicon probes with electrodes arranged as tetrodes (A4×2-tet configuration from NeuroNexus, Ann Arbor, MI). The shanks of the probes were marked with a fluorescent dye (DiI: Cat# V22885, or DiD: Cat# V22887, Thermo-Fisher Scientific) before penetration of the brain to assist in the identification of shank location postmortem. Before neural recordings, animals were head-fixed, the silicone elastomer was removed, and the electrodes were inserted through the craniotomy. The probe was held in a vertical position and lowered 2.9 mm from the brain surface. We waited for at least 15 minutes for the probe to settle before initiating recordings. Neural recordings were performed at multiple depths on each penetration, with recording sites typically 100-150 *µ*m apart to avoid recording from the same cells twice. Multiple penetrations were performed for each animal. A few recording sessions included only the presentation of one stimulus type (pure tones or AM noise), while the large majority of recording sessions included both.

A 50 *µ*m core diameter Polymicro optical fiber (Molex, part #1068001596) was attached to the silicon probe between the middle shanks and approximately 200 *µ*m above the top tetrode. The optical fiber was connected to a 445 nm laser calibrated to deliver 2 mW at the fiber tip.

### Estimation of recording location

At the conclusion of the experiments, animals were deeply anesthetized with euthasol and perfused through the heart with 4% paraformaldehyde. Brains were extracted and left in 4% paraformaldehyde for at least 24 hours before slicing. Brain slices (thickness 50 *µ*m or 100 *µ*m) were prepared under phosphate-buffered saline using a vibratome (Leica VT1000 S) and imaged using a fluorescence microscope (Axio Imager 2, Carl Zeiss) with a 1.25x and 2.5x objective (NA 0.16). To determine the locations of our recordings, we manually registered each histology slice containing dye fluorescence from a recording track to the corresponding coronal section in the Allen Mouse Common Coordinate Framework (Common Coordinate Framework v.3, © 2015 Allen Institute for Brain Science, Allen Brain Atlas API, available from http://brain-map.org/api/index.html). Recordings identified to be from the cerebral cortex were excluded from further analysis.

### Data analysis

#### Spike sorting and selection of D1-expressing neurons

Spiking activity was detected by applying a threshold (40-45 *µ*V) to bandpass (300 to 6000 Hz) filtered electrical signals measured by the electrodes. The activity from single units was isolated offline using the automated expectation maximization clustering algorithm Klustakwik (Kadir et al., 2014). Isolated clusters were only included in the analysis if less than 5% of inter-spike intervals were shorter than 2 ms. We also calculated a spike quality index, defined as the ratio between the peak amplitude of the spike waveform and the average variance, calculated using the channel with the largest amplitude. Cells were only included in the analysis if they had a spike quality index greater than 2. The analysis also excluded clusters identified as having noisy spike waveforms from visual inspection. Last, only neurons that had a firing rate (either spontaneous or evoked) of at least 1 spike/second were included in the analysis. Cells with lower firing rates were excluded because we considered our measurements from these potential cells to be unreliable.

Neurons were classified as D1-expressing if their onset response to laser stimulation (first 50 ms) was statistically larger (*p* < 0.01, Wilcoxon signed-rank test) than the baseline firing estimated from 200 ms before stimulus onset. Other neurons were classified as non-D1. Changing criteria slightly (*e*.*g*., requiring non-D1 neurons to have laser-evoked p-values greater than 0.1) did not qualitatively affect the results. Additional comparisons between neuron classes were performed while restricting the non-D1 population to only those neurons that were recorded from sites where D1 cells were found. That is, if no D1 neurons were observed on a tetrode during a session, cells from that tetrode were not included.

### Estimation of frequency tuning and responses to pure tones

To determine if a neuron was responsive to pure tones, we tested whether the evoked response during sound presentation (0 − 100 ms) was statistically different from the baseline spontaneous firing (measured during the 200 ms before sound onset) for any of the frequencies presented, collapsed across intensities. Because this test was performed for each of the 16 frequencies presented, we performed a Bonferroni correction for multiple comparisons (resulting in *α* = 0.05*/*16 = 0.0031). A tone response index (TRI) was calculated for the sound frequency that elicited the largest change from baseline firing for each neuron: TRI = (*r*_*e*_ −*r*_*b*_)*/*(*r*_*e*_ + *r*_*b*_), where *r*_*e*_ is the average evoked response for that sound frequency and *r*_*b*_ is the baseline spontaneous firing. To evaluate the sound frequency tuning of each neuron, we fit a Gaussian function to the average firing rate evoked by each frequency, collapsed across sound intensities. The frequency tuning bandwidth was estimated as the full width at half maximum of this function, which for a Gaussian corresponds to 2.355*σ*, where *σ* is the standard deviation. Only neurons with *R*^2^ values greater than 0.01 were included in the comparisons of tuning bandwidth. To estimate differences in response dynamics to pure tones, we calculated an onset-to-sustained index as OSI = (*r*_*o*_ −*r*_*s*_)*/*(*r*_*o*_ + *r*_*s*_), where *r*_*o*_ is the average firing rate during the early part of the stimulus (0 − 50 ms) and *r*_*s*_ is the average firing rate during the late part of the stimulus (50 − 100 ms).

### Estimation of responses to AM sounds

To determine if a neuron was responsive to amplitude modulated (AM) noise, we tested whether the evoked response was statistically different from the baseline spontaneous firing (measured during the 200 ms before sound onset) for any of the AM rates presented. Because neurons in the auditory system often show substantially different responses at the onset *vs*. the sustained periods of AM stimuli, we performed separate tests for each period: onset (0 − 100 ms) and sustained (100 − 500 ms). Because this test was performed for each of the 11 AM rates presented, we performed a Bonferroni correction for multiple comparisons (resulting in *α* = 0.05*/*11 = 0.0045). A sustained response index (SRI) was calculated for the amplitude modulation (AM) rate that elicited the largest change from baseline firing for each neuron: SRI = (*r*_*e*_ −*r*_*b*_)*/*(*r*_*e*_ + *r*_*b*_), where *r*_*e*_ is the average evoked response for that AM rate during the sustained period and *r*_*b*_ is the baseline spontaneous firing. AM rate selectivity was estimated by using an index that compared the maximum and minimum evoked firing rates during the sustained period across AM rates: AMRSI = (*r*_*max*_ − *r*_*min*_)*/*(*r*_*max*_ + *r*_*min*_). To estimate the synchronization of responses to AM sounds, we performed a Rayleigh’s test to determine the AM rates for which a neuron displayed statistically significant synchronization of its firing with respect to the phase of the amplitude modulation. This test included a Bonferroni correction for multiple comparisons (*α* = 0.0045). For each neuron, we estimated the highest AM rate where the neuron’s firing was significantly synchronized to the modulation period.

### Statistics

Throughout the study, we used non-parametric statistical tests implemented by the Python package SciPy (Virtanen et al., 2020). When comparing evoked firing rates to spontaneous rates (*e*.*g*., for responses to laser stimulation or sound stimulation), we used a test for related paired samples (Wilcoxon signed-rank test), where each trial provides one pair. When comparing measurements across two populations of cells, we used a non-parametric tests for two independent samples (Mann-Whitney U rank test). The comparisons associated with the histograms in Fig. 3D and Fig. 4E were performed using the absolute value of the response index in each case, while the triangles indicate the median values separately for positive and negative responses.

## Results

### Distinct classes of posterior striatal neurons respond to sound stimuli

To determine whether the representation of sounds differed across neuron classes in the posterior tail of the striatum, we recorded sound-evoked responses of photo-identified D1-expressing neurons and neighboring (non-D1) neurons in this brain region from naive awake mice using silicon multichannel probes that have electrodes organized as tetrodes (Fig. 1A). Photo-identification of D1 neurons during electrophysiological recordings was made possible by using DRD1::ChR2 mice which express the light-gated ion channel channelrhodopsin-2 (ChR2) in D1 neurons, and evaluating the spiking responses of each recorded neuron to laser stimulation delivered via an optical fiber attached to the recording probe. This method for in-vivo identification of genetically defined neuronal populations has been extensively used and validated in several brain regions, including the striatum (Lima et al., 2009; Kravitz et al., 2013; Lakunina et al., 2020). Figure 1B shows an example striatal neuron that responds reliably to laser stimulation; the quick strong response after laser onset indicates that this neuron expresses D1. This neuron also shows reliable responses to sound, a pure tone in this case (Fig. 1D). In the same brain region, we found neurons that showed no response to laser stimulation (Fig. 1C), but reliably respond to sound (Fig. 1E).

**Figure 1:**
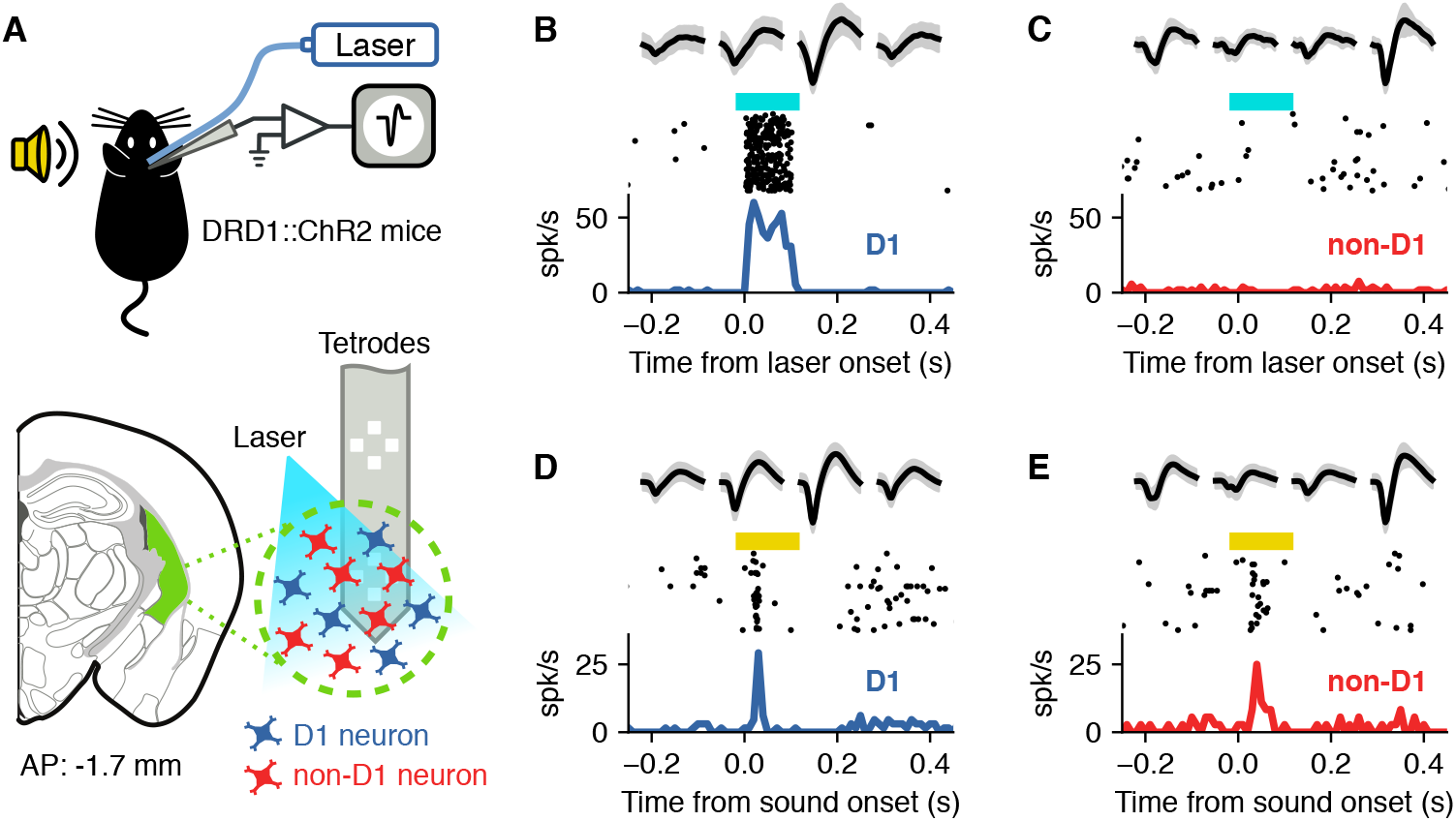
Photo-identification of D1-expressing striatal neurons. **A**, Extracellular recordings of neurons from the posterior tail of the striatum (green area) of awake head-fixed DRD1::ChR2 mice during sound presentation. D1-expressing neurons were identified during electrophysiological recordings by evaluating neural responses to blue laser stimulation. **B**, Example responses to laser stimulation from a striatal neuron. The top row shows the average (black) and standard deviation (gray) of the spike shape collected from each channel of a tetrode in the silicon probe. The middle row shows the firing for each presentation of the laser and the bottom row the peri-stimulus time histogram. The early and consistent response indicates this cell expresses ChR2 and therefore is a D1 neuron. **C**, Example of a cell that did not respond to the laser. Because of this, the cell is considered to be non-D1. **D**, Firing of the cell in panel B evoked by a 9.9 kHz pure tone. Note that the spike shapes match those in B. **E**. Firing of the cell in panel C evoked by a 9.9 kHz pure tone. Spike shapes match those in D.

From our sample of recorded cells, we identified 482 neurons as having statistically significant positive laser-evoked responses (Wilcoxon signed-rank test, *p* < 0.01) and therefore classified as D1-expressing, and 465 classified as non-D1 neurons. From these populations, we found that 43% of D1 neurons and 29% of non-D1 neurons showed a reliable evoked response to at least one sound in our stimulus ensemble, a mix of pure tones of different frequencies and amplitude modulated noise at different modulation rates. This difference in the fraction of responsive cells from each class was statistically significant (*p <* 0.0001, Fisher’s exact test), although is was much less apparent when the analysis was restricted to the 197 non-D1 cells that were recorded from sites where D1 cells were found (43% D1 *vs*. 40% non-D1 sound-responsive neurons, *p* = 0.027, Fisher’s exact test). Fig. 2 shows the estimated recording locations where we found sound-responsive neurons of each type (D1: blue, non-D1: red).

**Figure 2:**
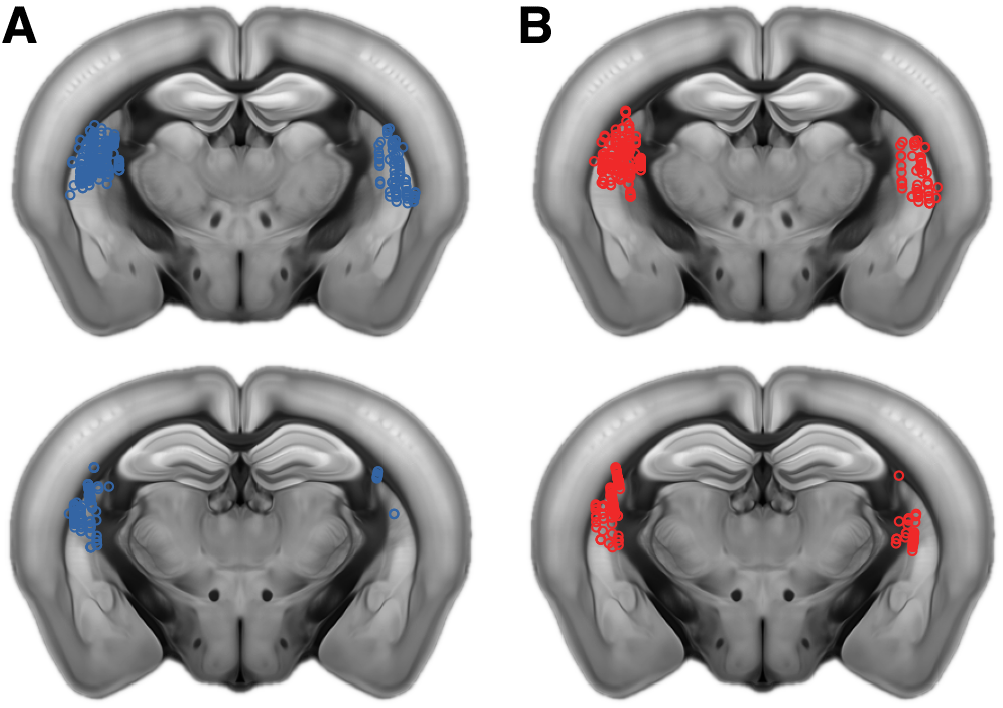
Location of sound-responsive striatal neurons. **A**, Coronal slices showing sites in both hemispheres where we found neurons that responded to at least one of the sounds in our ensemble (a combination of pure tones and amplitude modulated noise). Each circle represents one recording site. Sites are collapsed onto one of two anterior-posterior locations shown: approximately -1.35 mm (top) and -1.75 mm (bottom) from bregma. The ventral region of the posterior tail of the striatum was not sampled in our experiments. **B**, Coronal slices as in A showing sites where we found sound-responsive non-D1 neurons.

The observations described above indicate that subsets of medium spiny neurons in the posterior tail of the striatum that express the dopamine receptor D1, as well as neighboring non-D1 neurons, display reliable responses to sounds. We next wanted to test whether the sound-evoked responses and acoustic features encoded by these neurons differ between the two populations.

### D1-expressing neurons and their neighbors display similar sound frequency tuning

To test whether D1 neurons in the posterior striatum encoded the frequency of sounds with different fidelity compared to other neurons in this region, we evaluated the evoked responses of identified D1 and non-D1 cells to pure tones of different frequencies and intensities. A total of 400 D1 and 376 non-D1 neurons were recorded during the presentation of pure tones. Fig. 3A shows the average response of a D1 neuron for each frequency-intensity combination, indicating a clear tuning to specific frequencies and a dependence on sound intensity as is commonly observed across the auditory system. A subset of non-D1 neurons also displayed frequency selectivity (Fig. 3B), and both cell classes contained neurons in which the evoked response consisted of a decrease in firing, as illustrated with the example non-D1 cell in Fig. 3C.

**Figure 3:**
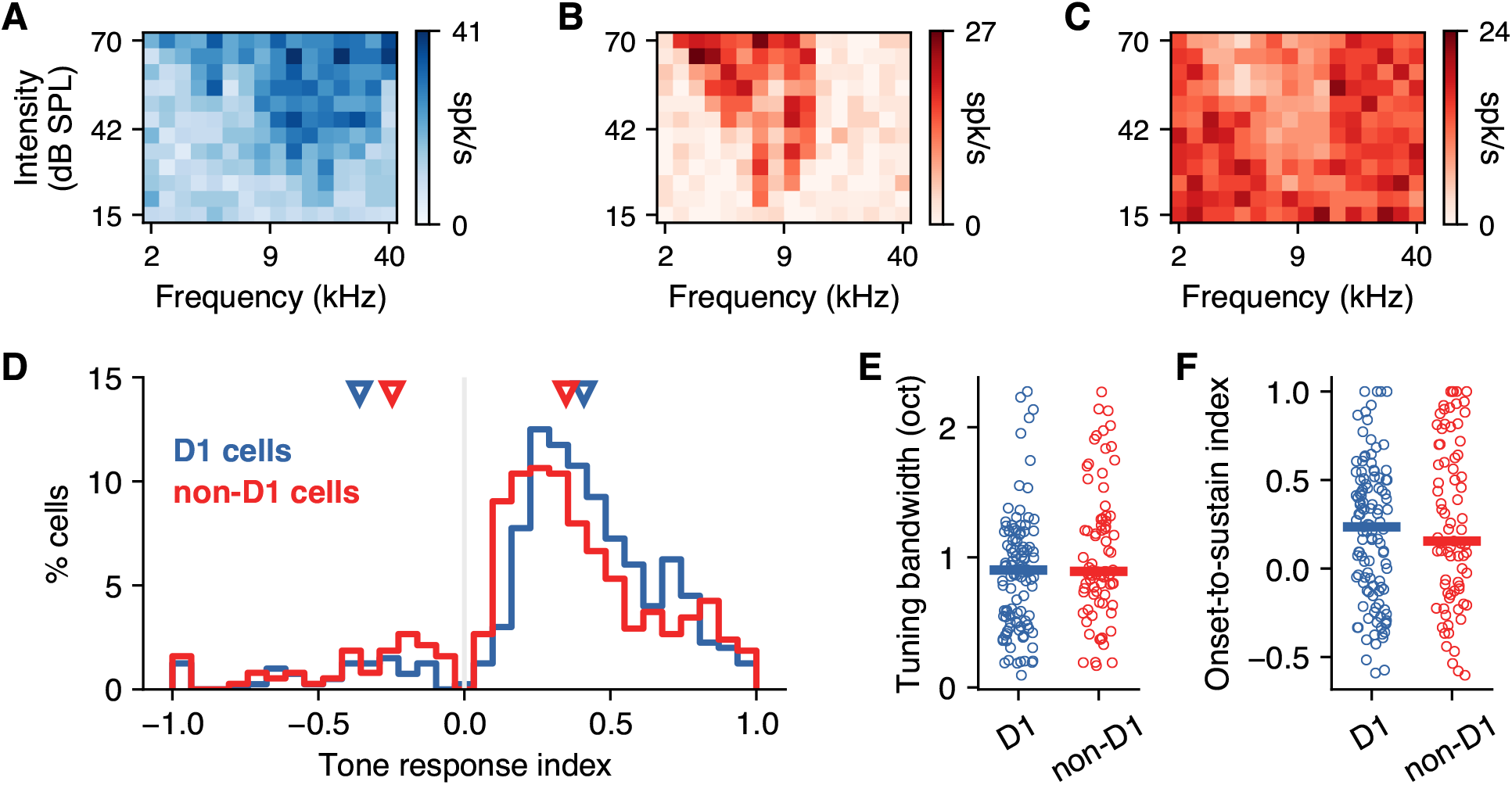
D1-expressing striatal neurons have comparable sound frequency tuning to neighboring neurons. **A**, Example frequency-intensity tuning curve from a D1 striatal neuron. Responses correspond to the average firing rate during the presentation of 100 ms pure tones. **B**, Example tuning curve from a neighboring non-D1 neuron. **C**, Example tuning curve from a non-D1 neuron that responded to sounds by decreasing its firing. **D**, Tone evoked-response index for the tone frequency that elicited the largest response in each neuron of each class. Many neurons responded by decreasing their firing. The triangles indicate the median response index calculated separately for neurons that responded by increasing or decreasing their firing. On average, D1 neurons showed larger responses than non-D1 neurons (*p* = 0.0006, Mann-Whitney U rank test). **E**, Frequency tuning bandwidth is similar between D1 and non-D1 neurons (*p* = 0.086, Mann-Whitney U rank test). This analysis includes only neurons for which a Gaussian curve can be fit to their tuning curve collapsed across intensities. Each circle corresponds to one neuron. Horizontal bars indicate the median. **F**, Firing dynamics (a comparison between onset and sustained firing) for the pure tone that elicited the strongest response is similar between D1 and non-D1 neurons (*p* = 0.507, Mann-Whitney U rank test).

We calculated a tone response index that compared the evoked response to the baseline firing for each neuron and plotted this index for the sound frequency that elicited the most reliable evoked response in each neuron (Fig. 3D). Positive values indicate that the evoked firing was larger than the spontaneous firing. As expected, most neurons (even if they did not have a statistically significant response to sounds) had firing rates evoked by the best stimulus that differed from the spontaneous firing, and therefore show a response index different from zero. Although these values largely overlapped across the two populations of cells, we found that the strongest sound-evoked changes in firing for D1 neurons were larger than those for non-D1 neurons (median absolute index: 0.41 for D1 *vs*. 0.35 for non-D1, *p* = 0.0006, Mann-Whitney U rank test). Moreover, the median values (illustrated with triangles in Fig. 3D) were larger for D1 neurons that had positive evoked responses and smaller for neurons with negative evoked responses, compared to median values for non-D1 neurons. This difference, however, disappeared when we restricted our analysis to the 188 non-D1 neurons recorded from sites where D1 cells were found (median: 0.42 for D1 *vs*. 0.44 for non-D1, *p* = 0.156, Mann-Whitney U rank test), suggesting that this result is influenced by the exact location of each recording site.

We next compared the tuning bandwidth of cells from each population. We first found cells for which responses across sound frequencies are well fit by a Gaussian function (115 out of 166 tone-responsive D1 neurons, and 79 out of 104 tone-responsive non-D1), and used the width at half maximum of this curve as an estimate of tuning bandwidth. We found no significant difference between the frequency tuning bandwidth across these populations of neurons (*p* = 0.086, Mann-Whitney U rank test). We then evaluated whether the dynamics of the responses to pure tones differed between D1 and non-D1 neurons. To this end, we estimated an index that compares the magnitude of the onset response (0 − 50 ms) to the sustained part of the response (50 − 100 ms) (Fig. 3F). We found no difference in this onset-to-sustained response index between the two populations (*p* = 0.507, Mann-Whitney U rank test).

These results suggest that responses to pure tones by D1-expressing neurons in the posterior striatum are comparable to those of their neighboring neurons. We next wanted to evaluate the encoding of temporal acoustic features by these neuronal populations.

### D1-expressing neurons have responses to AM sounds comparable to those of their neighbors

To test whether D1 neurons in the posterior striatum encoded temporal features of sounds with different fidelity compared to other neurons in this region, we evaluated the evoked responses of identified D1 and non-D1 cells to sinusoidally amplitude-modulated white noise at different modulation rates. A total of 475 D1 and 460 non-D1 neurons were recorded during the presentation of AM sounds. We found neurons from both classes that reliably responded to these sounds. As seen in neurons from other auditory regions, the responses of some cells were synchronized to the phase of the modulation, up to some modulation rate (Fig. 4A-D). While many neurons had the largest evoked firing for low modulation rates (Fig. 4A,C), other neurons were tuned to intermediate rates (Fig. 4B,D).

**Figure 4:**
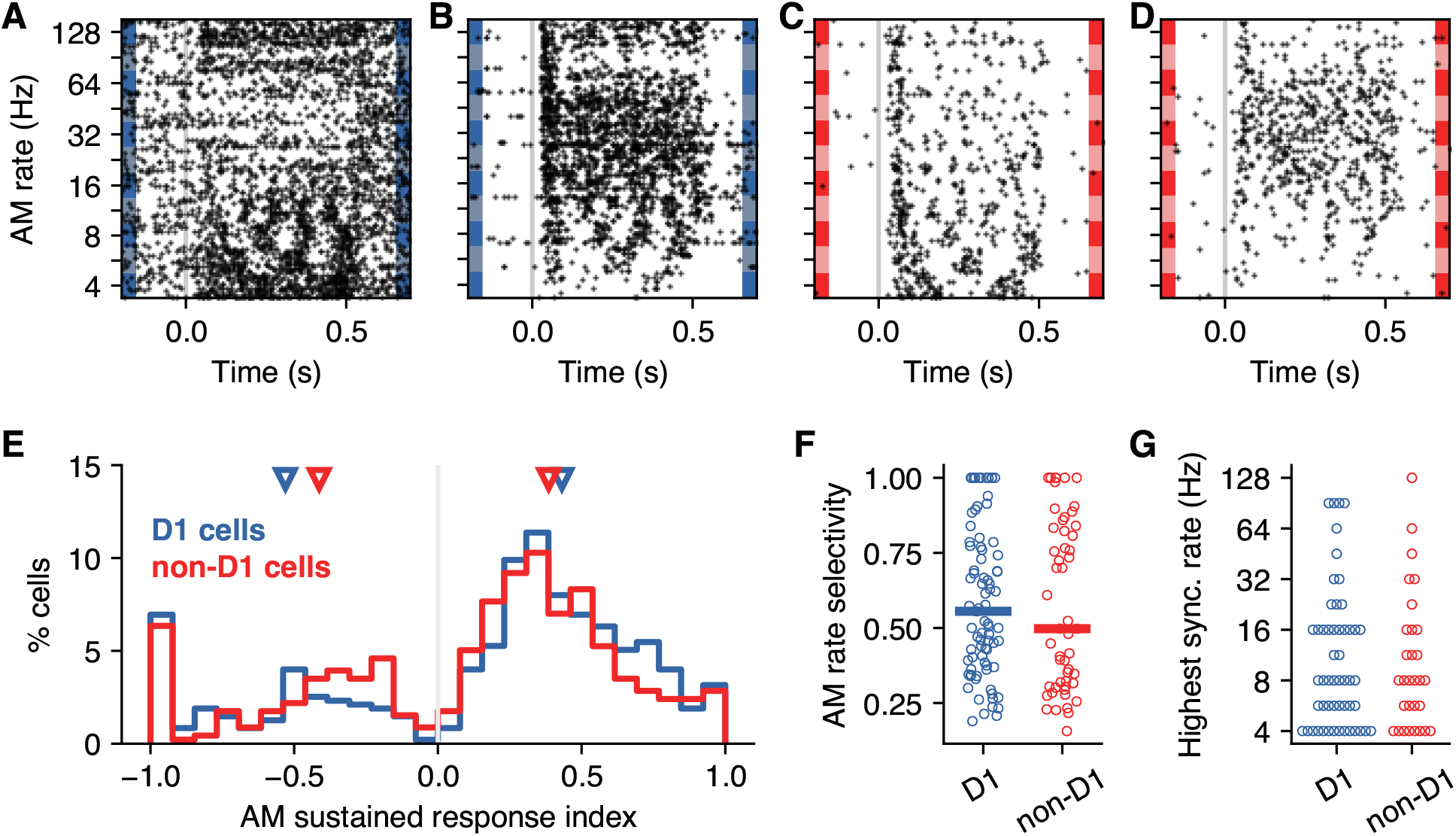
D1-expressing striatal neurons display similar selectivity to AM sound compared to their neighbors. **A**, Example responses of a D1 neuron to 500 ms amplitude-modulated (AM) white noise at different modulation rates. Synchronization of responses to the stimulus is clearly visible for low modulation rates. **B**, Responses to AM white noise for a different D1 neuron. This neuron is tuned to an intermediate modulation rate. **C**, Responses of a non-D1 neuron showing a high level of synchronization to the stimulus. **D**, Responses of a non-D1 neuron showing tuning to an intermediate modulation rate. **E**, AM sound evoked-response index for the stimulus that elicited the largest response in each neuron of each class. Responses are calculated for the sustain portion of the response (100-500 ms) Many neurons responded by decreasing their firing. The triangles indicate the median response index calculated separately for neurons that responded by increasing or decreasing their firing. On average, D1 neurons showed larger responses than non-D1 neurons (*p* = 0.0013, Mann-Whitney U rank test). **F**, AM rate selectivity is similar between D1 and non-D1 neurons (*p* = 0.578, Mann-Whitney U rank test). Selectivity was calculated by comparing the strongest to the weakest sustained response across AM rates. Only neurons with a statistically significant response to at least one AM rate are included. Each circle corresponds to one neuron. Horizontal bars indicate the median. **G**, The highest AM rate each neuron synchronized to was similar across cell classes (*p* = 0.463, Mann-Whitney U rank test).

We calculated a sustained response index that compared the evoked response to baseline firing for each neuron and plotted this index for the modulation rate that elicited the most reliable evoked response in each neuron (Fig. 4E). For many neurons, the firing rate during the sustained portion was lower than the spontaneous firing (indicated by negative values of the index). Although the values of this index largely overlapped across the two populations of cells, we found that the strongest sound-evoked changes in firing for D1 neurons were larger than those for non-D1 neurons (median absolute index: 0.47 for D1 *vs*. 0.39 for non-D1, *p* = 0.0013, Mann-Whitney U rank test). The median values (illustrated with triangles in Fig. 3D) were slightly larger for D1 neurons that had positive evoked responses and smaller for neurons with negative evoked responses, compared to median values for non-D1 neurons. This difference, however, disappeared when we restricted the analysis to non-D1 neurons recorded from sites where D1 cells were found (median: 0.47 for D1 *vs*. 0.49 for non-D1, *p* = 0.212, Mann-Whitney U rank test), suggesting that this result is influenced by the exact location of each recording site.

We next compared the modulation rate selectivity of cells from each population, estimated by comparing the maximum and minimum responses across modulation rates (including only neurons that were found to be responsive to AM sounds, 81 D1 and 52 non-D1 neurons). We found no significant difference between the AM rate selectivity across these populations of neurons (*p* = 0.832, Mann-Whitney U rank test). We then evaluated the highest modulation rate at which responses from each neuron synchronized to the stimulus (using the Rayleigh test for periodicity). We found no difference in the highest synchronized modulation rate between the two populations (*p* = 0.463, Mann-Whitney U rank test).

These results suggest that responses to amplitude modulated sounds by D1-expressing neurons in the posterior striatum are comparable to those of their neighboring neurons. Overall, the results above indicate that, in naive animals and outside the context of a task, different classes of neurons in the posterior tail of the striatum have access to and process acoustic features in a similar fashion.

## Discussion

In this study, we quantified the activity of neurons from the posterior tail of the striatum in response to sounds of different frequency or temporal structure. Optogenetic methods for identifying genetically distinct cell types during extracellular recordings allowed us to compare the representation of sounds by distinct classes of striatal neurons. Specifically, we focused on one of the major cell classes in the posterior striatum, namely those that express the dopamine receptor D1 (and form the direct striatonigral pathway) and compared them to other neurons in the same striatal region. The cellular composition of the striatum is such that D1-expressing and D2-expressing neurons are present in similar proportions and together account for more than 90% of striatal neurons (Kreitzer and Malenka, 2008). Therefore, it is likely that most of our recorded non-D1 neurons are in fact D2-expressing indirect-pathway neurons. The possibility of observing differences in the representation of sounds by these cell classes was motivated by previous studies that suggest differential innervation of striatal pathways by cortical neurons (Lei et al., 2004; Kreitzer and Malenka, 2007; Wall et al., 2013). We found, however, that the representation of spectral and temporal features of sounds by D1-expressing neurons is similar to the representation of these features by neighboring neurons.

A common concern when applying methods for photo-identification during electrophysiological recordings is the possibility of observing laser-evoked responses from multisynaptic indirect activation by other neurons. This concern is less relevant when identifying striatal D1-expressing neurons as they are GABAergic in nature (and therefore their activation will result in inhibition of their synaptic partners) and minimized by using only the early portion of the laser-evoked responses for identification (before recurrence can have a major impact).

Our results indicating that sound-responsive D1 neurons encode acoustic features with similar fidelity to that of sound-responsive non-D1 neurons was supported by multiple measures and it was robust to applying stricter criteria for inclusion of cells (*e*.*g*., using only cells with high firing rates to increase the reliability of response estimates; data not shown). However, comparisons across cell classes suggested different results when we included all identified neurons or included only neurons recorded on the same electrodes. These observations cast doubt on the validity of the observed differences across cell classes, as these effects could be explained by distinct levels of responsiveness by cells from different recorded locations. Variability in the recording locations across mice, together with the limited precision of our method for estimating the location of each recording make it impractical to derive further conclusions from our data regarding these potential differences.

Previous studies have observed strong selectivity to sound frequency in the responses of posterior striatal neurons of mice trained to perform auditory tasks (Guo et al., 2018; Chen et al., 2019). Our study complements these observations by demonstrating that these neurons display robust responses to sounds even in naive mice. Moreover, our study illustrates that the responses of posterior striatal neurons can be synchronized to the amplitude modulation of the stimulus or be tuned to specific modulation rates, as observed in auditory thalamic and cortical neurons (Ponvert and Jaramillo, 2019). One of these earlier studies evaluated the evoked responses to pure tones from different classes of striatal neurons identified according to their spike shapes (Chen et al., 2019). They found that subsets of neurons from all identified classes (medium spiny neurons, cholinergic interneurons, and fast-spiking interneurons) displayed responses to tones with various dynamics and frequency tuning. Because the group of cells classified as medium spiny neurons contains both direct and indirect pathway neurons, their study could not derive conclusions regarding potential differences between cells from these two pathways. Our study complements these results by demonstrating that direct pathway neurons encode sounds in a similar fashion to their neighbors. Because an overwhelming majority of neurons in the striatum are projection medium spiny cells, split evenly between the two striatal output pathways (Kreitzer and Malenka, 2008), it is likely that this conclusion extends to the comparison between direct *vs*. indirect pathway neurons.

These observations indicate that multiple classes of posterior striatal neurons have access to detailed spectro-temporal acoustic features and could therefore potentially influence behavioral responses to sounds. It remains unknown whether non-acoustic features that influence the activity of posterior striatal neurons, such as reward expectation and choice-related variables (Guo et al., 2019), are also represented similarly across cell classes, or whether differences in the representation of sounds emerge when investigating subgroups of neurons within each class (Gokce et al., 2016).

## Acknowledgements

This research was supported by the National Institute on Deafness and Other Communication Disorders (R01DC015531), and the Office of the Vice President for Research & Innovation at the University of Oregon.

